# Generation of tumorigenic porcine pancreatic ductal epithelial cells: toward a large animal model of pancreatic cancer

**DOI:** 10.1101/267112

**Authors:** Neeley Remmers, Jesse L. Cox, James A. Grunkemeyer, Shruthi Aravind, Christopher K. Arkfeld, Michael A. Hollingsworth, Mark A. Carlson

## Abstract

**Background**. A large animal model of pancreatic cancer would permit development of diagnostic and interventional technologies not possible in murine models, and also would provide a more biologically-relevant platform for penultimate testing of novel therapies, prior to human testing. Here, we describe our initial studies in the development of an autochthonous, genetically-defined, large animal model of pancreatic cancer, using immunocompetent pigs.

**Methods**. Primary pancreatic epithelial cells were isolated from pancreatic duct of domestic pigs; epithelial origin was confirmed with immunohistochemistry. Three transformed cell lines subsequently were generated from these primary cells using expression of oncogenic KRAS and dominant negative p53, with/without knockdown of p16 and SMAD4. We tested these cell lines using *in vitro* and *in vivo* assays of transformation and tumorigenesis.

**Results**. The transformed cell lines outperformed the primary cells in terms proliferation, population doubling time, soft agar growth, 2D migration, and Matrigel invasion, with the greatest differences observed when all four genes (KRAS, p53, p16, and SMAD4) were targeted. All three transformed cell lines grew tumors when injected subcutaneously in nude mice, demonstrating undifferentiated morphology, mild desmoplasia, and staining for both epithelial and mesenchymal markers. Injection into the pancreas of nude mice resulted in distant metastases, particularly when all four genes were targeted.

**Conclusions**. Tumorigenic porcine pancreatic cell lines were generated. Inclusion of four genetic “hits” (KRAS, p53, p16, and SMAD4) appeared to produce the best results in our *in vitro* and *in vivo* assays. The next step will be to perform autologous or syngeneic implantation of these cell lines into the pancreas of immunocompetent pigs. We believe that the resultant large animal model of pancreatic cancer could supplement existing murine models, thus improving preclinical research on diagnostic, interventional, and therapeutic technologies.

## Introduction

In the United States in 2016, approximately 53,000 people (48% female) were diagnosed with pancreatic cancer (~3.1% of all new cancer diagnoses), and there were ~42,000 deaths (49% female) from pancreatic cancer (~7.0% of all cancer deaths) [*1,2*]. The lifetime risk for pancreatic cancer is approximately 1 in 65 [*1,2*]. The incidence of pancreatic cancer has been gradually increasing since the mid-1990’s, and generally is higher in the African-American population [*1,2*]. Pancreatic cancer is now is the fourth most common cause of cancer-related death in both men and women (after lung, prostate, and colorectal cancer, or lung, breast, and colorectal cancer, respectively) [*1,2*]. Despite apparent advances in treatment modalities and strategies [*3*], mortality from pancreatic cancer has not decreased [*1,2*]. As of 2012, the U.S. overall 5-year survival rate from pancreatic cancer was 7.7%; 5-year survival rates in localized, regional (nodal spread), or metastatic disease were 29.3, 11.1, and 2.6%, respectively [*1,2*]. So there remains a need for improved early diagnosis and therapy for pancreatic cancer.

Rodent models of pancreatic cancer may not accurately reflect human biology because of differences in physiology, anatomy, immune response, and genetic sequence between the two species [*4–7*]. Remarkably, only 5–8% of anti-cancer drugs that emerged from preclinical studies and entered clinical studies have been ultimately approved for clinical use [*8,9*]. The cause of this low approval rate is multifactorial, but likely includes the less-than-optimal predictive ability of some murine models (e.g., tumor xenografting into immunosuppressed mice) to determine the efficacy of various therapeutics in humans [*4–6,10–14*]. Moreover, there are a number of genes for which the genotype-phenotype relationship is discordant between mice and human, including *CFTR^-/-^* and *APC*^+/-^ [*15,16*]. Incidentally, both the porcine *CFTR*^-/-^ and *APC*^+/-^ mutants reiterate the human phenotype (pulmonary/GI disease and rectal polyposis, respectively) [*15–17*], in contradistinction to the murine mutants.

In fairness, the recent trend to employ genetically-engineered mouse models (GEMM), patient-derived xenografts (PDX), humanized mice, and *in vivo* site-directed CRISPR/Cas9 gene-edited mice in the testing of anti-cancer therapeutics may yield murine models with better predictive ability than obtained with previously [*6,18–22*]. Though promising, these more advanced murine models come with increased cost and complexity [*20*], and experience with them still is early. Importantly, all murine models have limited utility in the development of diagnostic or interventional technology that requires an animal subject whose size approximates a human. So at present, there remains a need for improved animal models of pancreatic cancer that (1) are more predictive of human response to anti-cancer therapy [*20,22*], and (2) are of adequate size for development of specific technologies. Herein we describe some initial steps taken in the development of a genetically-defined, autochthonous model of pancreatic cancer in immunocompetent pigs.

## Materials and Methods

### Standards, rigor, reproducibility, and transparency

The animal studies of this report were designed, performed, and reported in accordance with both the ARRIVE recommendations (Animal Research: Reporting of *In Vivo* Experiments [*23*]) and the National Institutes of Health Principles and Guidelines for Reporting Preclinical Research [*24,25*]; for details, refer to Tables S1 and S2, respectively.

### Materials and animal subjects

All reagents were purchased through Thermo Fisher Scientific (www.thermofisher.com) unless otherwise noted. Short DNA sequences for vector construction, mutagenesis, and amplification purposes are shown in Table S3. Antibody information is given in Table S4. Wild type domestic swine (male and female; age 3 months at time of purchase; 30–32 kg) were purchased from the Animal Research and Development Center of the University of Nebraska Lincoln (ardc.unl.edu). Athymic homozygous nude mice (Crl:NU(NCr)-*Foxn*1*^nu^*; female; 8–9 weeks old) were purchased from Charles River Laboratories, Inc. (www.criver.com). Primers utilized in this report (Table S3) were synthesized by Integrated DNA Technologies, Inc. (www.idtdna.com). DNA sequencing was performed by the UNMC Genomics Core Facility (www.unmc.edu/vcr/cores/vcr-cores/genomics). Oncopigs [*26*] were purchased from the National Swine Resource and Research Center (NSRRC; www.nsrrc.missouri.edu).

### Animal welfare

The animals utilized to generate data for this report were maintained and treated in accordance with the *Guide for the Care and Use of Laboratory Animals* (8^th^ ed.) from the National Research Council and the National Institutes of Health [*27*], and also in accordance with the Animal Welfare Act of the United States (U.S. Code 7, Sections 2131 – 2159). The animal protocols pertaining to this manuscript were approved by the Institutional Animal Care and Use Committee (IACUC) of the VA Nebraska-Western Iowa Health Care System (ID numbers 00927, 00937, 00998, and 01017) or by the IACUC of the University of Nebraska Medical Center (ID number 16–133–11-FC). All procedures were performed in animal facilities approved by the Association for Assessment and Accreditation of Laboratory Animal Care International (AAALAC; www.aaalac.org) and by the Office of Laboratory Animal Welfare of the Public Health Service (grants.nih.gov/grants/olaw/olaw.htm). All surgical procedures were performed under isoflurane anesthesia, and all efforts were made to minimize suffering. Euthanasia was performed in accordance with the AVMA Guidelines for the Euthanasia of Animals [*28*].

### Porcine operative procedures

Further details on transgenic porcine subjects and related welfare, safety, husbandry, operative procedures, and perioperative management are given in the Supporting Information.

### Isolation of porcine pancreatic ductal epithelial cells

A detailed protocol for isolation of porcine pancreatic ductal epithelial cells is provided in the Supporting Information (**Protocol S1**). In brief, the intact pancreas from male porcine research subjects (age 5 mo) was harvested within 5 min after euthanasia, which was accomplished by transection of the intrathoracic inferior vena cava and exsanguination while under deep isoflurane anesthesia. These pigs had been on a protocol to study biomaterials within skin wounds of the dorsum. The subject had not received any recent medication other the anesthetics given for euthanasia; buprenorphine and cefovecin sodium had been given 4 weeks prior to euthanasia. Immediately after explantation of the pancreas, the main pancreatic duct was dissected sterilely with micro instruments from the organ body under 3.5x loupe magnification. The duct then was mechanically digested by passage through a 70 μm sieve (Corning™ Sterile Cell Strainers, Thermo Fisher Scientific, cat. no. 07–201–431).

The collected fragments were enzymatically digested with 1 mg/mL of Collagenase D at 37°C for 1 h with gentle shaking. The cells were pelleted (600 *g* × 5 min), the supernatant was discarded, the cell pellet was resuspended in whole media, which was defined as: DMEM (high glucose with L-glutamine; Thermo Fisher Scientific, cat. no. 12100–046) supplemented with 10% (final concentration) fetal bovine serum (FBS; Thermo Fisher Scientific, cat. no. 26140079) and 1% Antibiotic-Antimycotic Solution (Corning Inc., cat. no. 30–004-CI; cellgro.com). Cell concentration in the resuspension was determined with a hemocytometer, and cells then were diluted and pipetted into a 96-well plate (1–10 cell/well, 100–200 μL/well). After 5–7 days of culture under standard conditions (whole media, 37°C, 5% CO_2_), wells that contained cells with epithelial-like morphology were trypsinized and re-plated into a new 96-well plate, in order to dilute out any fibroblasts. Cells were passaged in this fashion at least four times, until no cells with fibroblast morphology were present. The resulting cells were passaged up to a T25 flask, and maintained with standard conditions.

### Generation of p53 and KRAS mutants and construction of expression vector

In order to generate the porcine p53^R167H^ mutant, wild-type p53 cDNA first was amplified from cervical lymph node tissue, which was obtained <5 min after euthanasia of a 4-month-old male domestic swine that had been on an unrelated research protocol. In brief, fresh nodal tissue was flash-frozen in liquid N2 and then pulverized with a mortar and pestle, with continual addition of liquid N2 during pulverization. The frozen powder then was placed into the first buffer solution of the QIAGEN RNEasy Mini Kit (cat. no. 74104; www.qiagen.com), and total RNA was isolated per the manufacturer’s instructions.

After isolation, the total RNA underwent reverse transcription to cDNA with a Verso cDNA Synthesis Kit (Thermo Fisher Scientific, cat. no. AB1453A), per the manufacturer’s instructions. The wild type p53 sequence was amplified out of the cDNA using the PCR primers shown in Table S3, which flanked the p53 cDNA with *Sal*I and Bam*HI* restriction sites. Successful amplification of the wild-type p53 cDNA was confirmed by inserting the amplified candidate sequence into the TOPO^®^ vector (TOPO^®^ TA Cloning^®^ Kit; Invitrogen™/Life Technologies™, Thermo Fisher Scientific, cat. no. K202020) per the manufacturer’s instructions, followed by sequencing.

Site-directed mutation of wild-type p53 into p53^R167H^ was performed using Agilent Technologies’ QuickChange II Site-Directed Mutagenesis Kit (cat. no. 200523; www.genomics.agilent.com) with the mutagenic primers shown in Table S3, per the manufacturer’s instructions. Presence of the p53^R167H^ mutation was verified by sequencing as described above. The multiple cloning site of a pIRES2-AcGFP1 Vector (Takara Bio USA, Inc., cat. no. 632435; www.clontech.com; manufacturer’s vector information included as **Fig. S1**) was cut with *Sal*I and Bam*HI*, and the p53^R167H^ sequence then was ligated into this plasmid.

The source of the porcine KRAS^G12D^ mutant was the plasmid used to generate the p53/KRAS Oncopig [*26,29*]. The KRAS^G12D^ cDNA was amplified out of this plasmid with primers (see Table S3) that flanked the sequence with *Xho*I and *Pst*I restriction sites. The amplified product was inserted into the TOPO vector and verified by sequencing, as described above. The above pIRES2-AcGFP1 Vector (already containing the p53^R167H^ sequence) then was cut with *Xho*I and *Pst*I, and the KRAS^G12D^ sequence was ligated into this plasmid, producing a pIRES2-AcGFP1 Vector which contained both mutant cDNAs within its multiple cloning site (KRAS^G12D^ upstream).

The newly-constructed plasmid, hereafter designated as GKP (G = AcGFP1; K = KRAS^G12D^; P = p53^R167H^), was transformed into One Shot™ Stbl3™ Chemically Competent *E. coli* (Invitrogen™/Thermo Fisher Scientific, cat. no. C737303), per the manufacturer’s instructions, and plasmid DNA subsequently was isolated using a QIAGEN Plasmid Maxi Kit (cat. no. 12162), per the manufacturer’s instructions. This plasmid then was transfected into Takara’s Lenti-X™ 293T cells (Clontech, cat. no. 632180), using Takara’s Xfect™ Transfection Reagent (Clontech, cat. no. 631317), per the manufacturer’s instructions, to generate infectious lentiviral particles that would direct expression of AcGFP1, KRAS^G12D^, and p53^R167H^ mutants in transduced cells.

### Generation of shRNA-expressing vectors

Short hairpin RNA (shRNA) constructs targeting the porcine SMAD4 and p16 genes were created using InvivoGen’s siRNA Wizard™ software (www.invivogen.com/sirnawizard). Three targeting sequences for each gene along with scrambled controls initially were generated and tested. The shRNA construct that demonstrated the best target knock-down in preliminary experimentation (as verified by PCR, data not shown) was utilized for subsequent experiments (see Table S3 for sequences ultimately selected for the shRNA constructs). Primers were cloned into the psiRNA-h7SKhygro G1 and psiRNA-h7SKneo G1 vectors (InvivoGen, cat. no. ksirna3-h21 and ksirna3-n21, respectively; www.invivogen.com) and plasmids then were isolated, all per the manufacturer’s protocol.

### Cell transformations

Primary porcine pancreatic epithelial cells in T75 flasks were grown to 80% confluency under standard conditions. The media then was exchanged with 2–3 mL of supernatant from non-lysed Lenti-X™ 293T cells (containing GKP viral particles) with 2 μg/mL polybrene (cat. no. TR1003, Thermo Fisher Scientific). After 24–48 h at 37°C, treated epithelial cells were re-seeded into 6-well plates under standard conditions and grown to 80% confluency. An exchange with whole media containing 2 μg/mL G418 aminoglycoside antibiotic then was performed; the G418 dose was chosen based on preliminary dose-response studies against non-treated epithelial cells. After 24 h, a whole media exchange was done, and the presence of transduced cells was determined with inverted GFP fluorescent microscopy of living cells. Subsequent transfections for RNAi were done with the above plasmids employing shRNA sequences against SMAD4 and/or p16, and using the LyoVec™ reagent (InvivoGen, cat. no. lyec-12), all per the manufacturer’s protocol. Transfected cells then were selected for expression of the shRNA vector using the appropriate aminoglycoside antibiotic (G418 or hygromycin B).

### PCR

Cell and tissue RNA was isolated using the QIAGEN RNEasy Mini Kit. Purified RNA then was used to generate cDNA using the Verso cDNA Synthesis Kit. The Platinum^®^ Blue PCR Supermix (Invitrogen™/Life Technologies, cat. no. 12580) subsequently was used for all PCR reactions. Amplified products were separated with agarose gel electrophoresis, and then visualized using a UV-light box. qPCR was performed using the PowerUp™ SYBR^®^ Green Master Mix (Applied Biosystems™/ Thermo Fisher Scientific, cat. no. A25741) per manufacturer’s protocol, and run on an Applied Biosystems™ 7500 Fast Dx Real-Time PCR Instrument. Fold changes in gene expression were calculated using the comparative *C*_T_ method [*30*]. All primers used are listed in Table S3.

### Immunoblotting

Western blot analysis was performed to confirm overexpression of the mutant p53 protein (see Table S4 for a list of antibodies used), as previously described [*31*]. An antibody specific for the mutant KRAS protein was not commercially available. Antibody expression was visualized using the Li-Cor Odyssey Electrophoresis Imaging System (www.licor.com).

### Soft agar assay

A standard soft agar assay [*32*] was used to determine anchorage independent growth. A base layer of 1% agarose was plated into 6-well plates. A total of 2,500 cells/well were mixed with 0.7% agarose and plated on top of the base layer. The plates were incubated under standard conditions for 21 days. The cells then were stained with crystal violet, and counted using an inverted microscope. Cells were plated in triplicate, and total counts from all three wells were averaged.

### Migration assay

A standard scratch assay (monolayer wounding) [*33*] was performed to determine cellular migration rate. Cells were plated in triplicate into 6-well plates. A horizontal scratch using a 10 μL pipet tip was made in each well. After washing away scratched-off cells, baseline images along the scratch were obtained, the plates were incubated under standard conditions, and subsequent images were captured at 3, 6, 9, 12, and 15 h after the initial scratch. ImageJ software (imagej.nih.gov/ij) was used to measure the distance between the two migrating cellular fronts (scratch edges) at 3–5 locations along the scratch. Average distance at each time point was plotted to generate the migratory rate (μm/h).

### Invasion assay

BioCoat™ Matrigel™ Invasion Chambers (Corning™, Thermo Fisher Scientific, cat. no. 08–774) were plated with 50,000 cells (upper chamber) in triplicate, and incubated under standard conditions for 24 h. The media from the upper chamber then was removed, and any cells remaining in the upper chamber were removed using a cotton swab. Cells that had migrated to the bottom of the membrane were stained using a Kwik-Diff™ kit (Shandon™, Thermo Fisher Scientific, cat. no. 9990701). Membranes were mounted onto glass slides, and cells per high-power field were counted using ImageJ software.

### Population doubling assay

Cells were plated in 6-well plates (20,000 cell/well), and cultured under standard conditions. Triplicate plates then were trypsinized on days 1, 2, 3, 4, 6, and 8, and cells were counted with a hemocytometer. Cell number *vs*. day was plotted to determine the day range in which linear growth was achieved. The data from this linear growth phase were used to determine population doubling time (DT) using the formula: DT = (Δt) × ln(2) ÷ ln(N*_f_*/N*_i_*) where Δt = time interval between initial and final cell count, N*_f_* = cell count at final time, and N*_i_* = cell count at initial time.

### Proliferation assay

Relative cell proliferation rates were assayed using an MTT (3-(4,5-dimethylthiazol-2-yl)-2,5-diphenyltetrazolium bromide) assay kit (Vybrant™ MTT Cell Proliferation Assay Kit, Invitrogen™, Thermo Fisher Scientific, cat. no. V13154). Cells were plated in triplicate in a 96-well plate (5,000 cell/well), and cultured under standard conditions for 48 h. MTT reagent then was added to the cells per the manufacturer’s instructions, followed by addition of the solvent solution 3.5 h later. Absorbance was measured with a plate reader 3.5 h after solvent addition. Mean absorbance was normalized to absorbance from wild type pancreatic ductal epithelial cells to calculate fold-difference in proliferation.

### Immunofluorescence and immunohistochemistry

Antibodies used in immunofluorescent and immunohistochemical experiments are listed in Table S4. Agilent Dako EnVision kits (www.agilent.com) were used for all IHC analyses per the manufacturer’s instructions.

### Subcutaneous tumorigenic cell injection

Subcutaneous implantation of tumorigenic cells was performed as previously described [*34*], with some modifications. Transformed porcine pancreatic ductal epithelial cells (the three lineages described in Table 1) were trypsinized, counted, and resuspended in DMEM at a concentration of 1 × 10^7^ viable cells/mL. Nude mice (N = 30; 100% female; maintained in microisolator cages with soft bedding and fed regular chow) were randomized into three treatment groups (representing each transformed cell line in Table 1; N = 10 mice per group, 100% female) using an online randomization tool. Mice then were injected with 5 × 10^6^ cells (500 μL) into the right hind flank under brief isoflurane inhalational anesthesia, administered with a Matrx VMS^®^ small animal anesthesia machine, within a small animal operating room. Tumors were allowed to grow for 6 weeks or until they reached 2 cm in diameter, as measured with a caliper, and then subjects were euthanized using an AVMA-approved [*28*] method of CO_2_ asphyxiation. At necropsy all gross tumor was measured and collected, portions underwent formalin fixation and paraffin embedding, and sections subsequently underwent H&E or immunohistochemical staining as described above. An independent, a blinded pathologist analyzed the stained sections to determine whether tumors were epithelial in origin, and if they displayed malignant features.

**Table 1.**
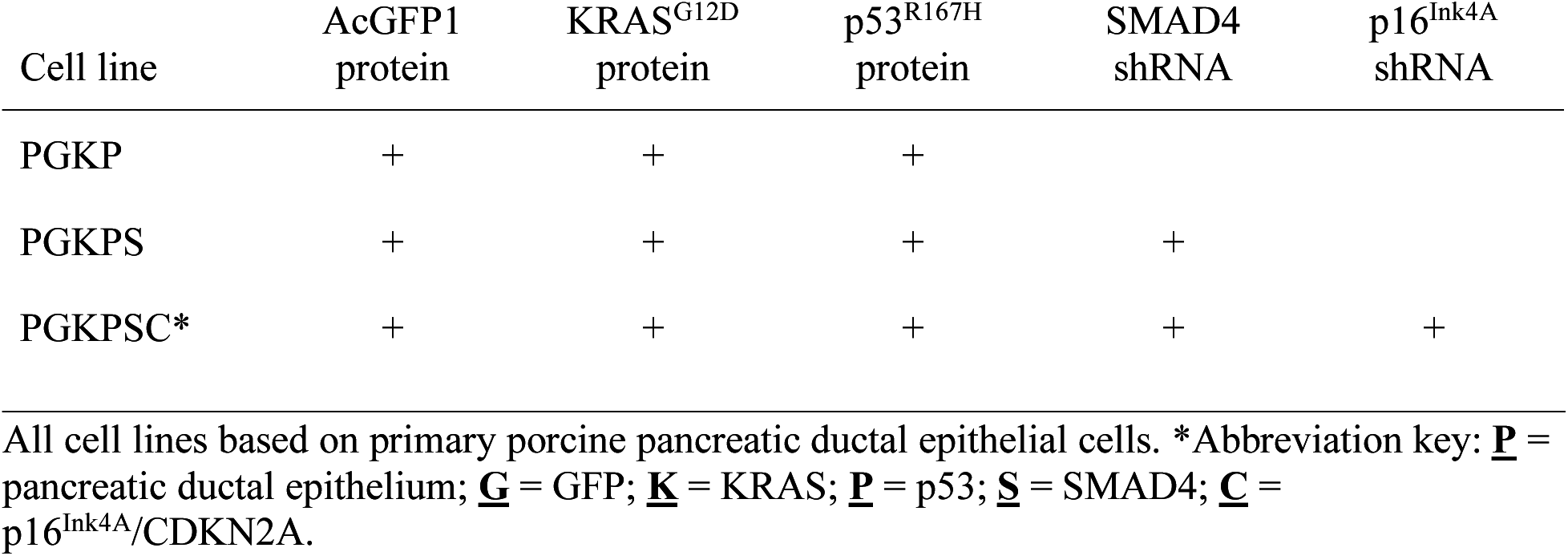
Tumorigenic cell lines and their expression products.

| Cell line | AcGFP1<br>protein | KRAS<sup>G12D</sup><br>protein | p53<sup>R167H</sup><br>protein | SMAD4<br>shRNA | p16<sup>Ink4A</sup><br>shRNA |
| --- | --- | --- | --- | --- | --- |
| PGKP | + | + | + |  |  |
| PGKPS | + | + | + | + |  |
| PGKPSC* | + | + | + | + | + |

All cell lines based on primary porcine pancreatic ductal epithelial cells. \*Abbreviation key: **P** = pancreatic ductal epithelium; **G** = GFP; **K** = KRAS; **P** = p53; **S** = SMAD4; **C** = p16<sup>Ink4A</sup>/CDKN2A.

### Orthotopic tumorigenic cell injection

Orthotopic implantation of tumorigenic cells was performed as previously described [*34*] to analyze metastases and desmoplasia. In brief, transformed porcine pancreatic ductal epithelial cells (the three lineages described in Table 1) were trypsinized and counted, and 1 × 10^4^ viable cells were suspended into 20 μL DMEM. Nude mice (N = 36; 100% female) housed as described above were randomized into three treatment groups (representing each transformed cell line in Table 1; N = 12 per group, 100% female) using an online randomization tool. The 20 μL cell suspension then was injected with a 20-gauge needle into the pancreas of each nude mouse through a 5 mm incision in the left upper quadrant, under isoflurane anesthesia within a small animal operating room. Mice were euthanized 6 weeks after injection using CO_2_ asphyxiation as described above, and tumors and organs were harvested for gross and histologic analysis, as described in the previous paragraph.

### Statistics and power analysis

Data are reported as mean ± standard deviation. Groups of continuous data were compared with ANOVA and the unpaired t-test. Categorical data were compared with the Fisher or Chi square test. For the power analysis of the murine subcutaneous tumor implant assay, tumor diameter was selected as the endpoint. Setting alpha = 0.05 and power = 0.8, ten mice per group were needed across three groups to detect a difference in means of 30% with the standard deviation estimated at 20% of the mean. In the orthotopic implantation assay, N = 10 mice per treatment group across three treatment groups were needed to detect a 100% difference in effect (+tumor) at a single metastatic site (with alpha set at 0.05 and power = 0.8); or, combining all seven metastatic sites together, N = 10 mice per group were needed to detect a 40% difference in effect.

## Results

### Isolation of primary porcine pancreatic ductal epithelial cells

Cells cultured from micro-dissected pancreatic ducts displayed epithelial morphology under phase microscopy and stained for CK19 (an established marker of pancreatic ductal epithelium [*35*]; **Fig. 1A-B**). Based on these results, we were confident that we had a population of pancreatic epithelial cells that we could use to generate tumorigenic cell lines.

**Fig. 1.**
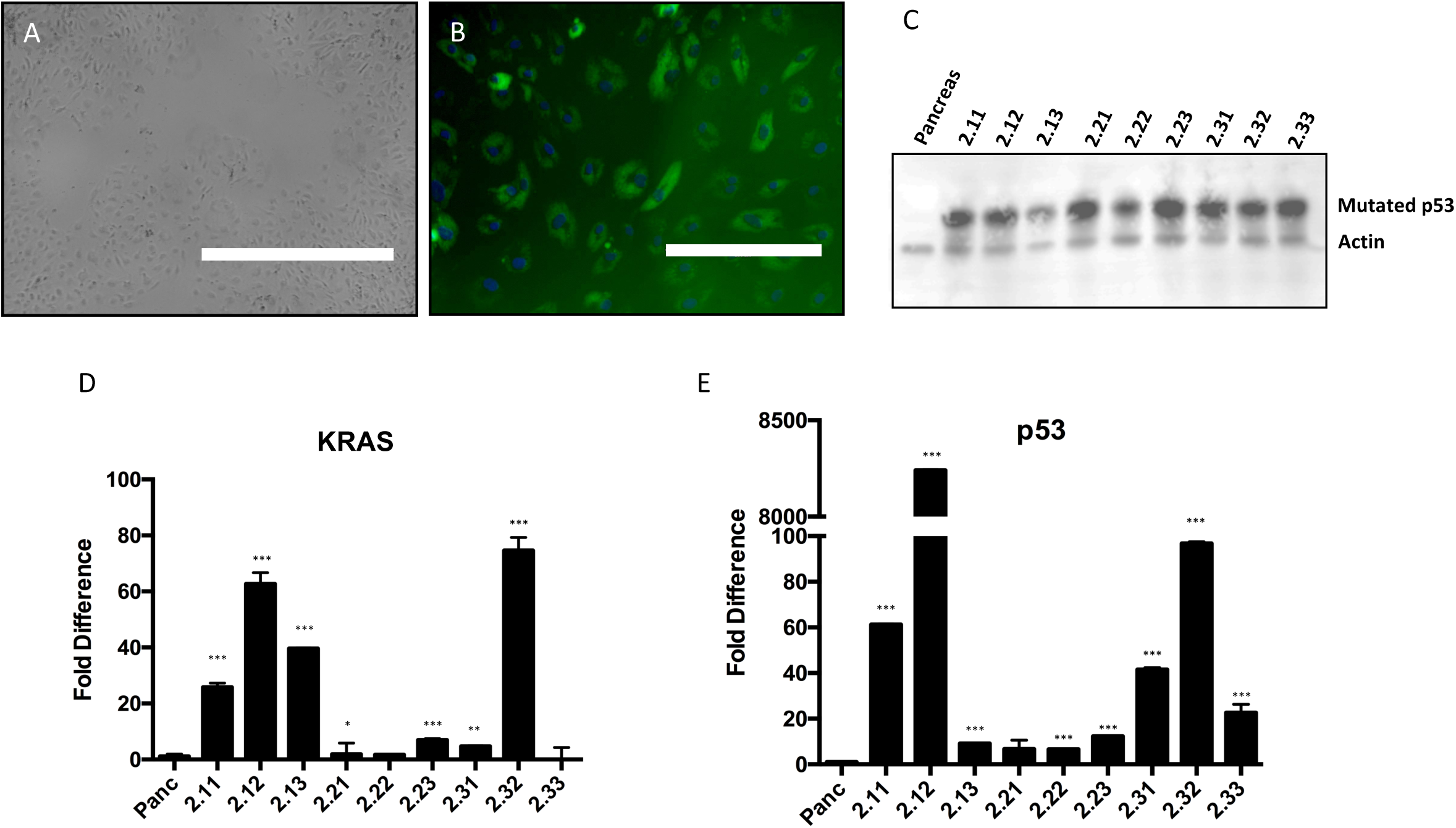
Isolation and transduction of primary porcine pancreatic ductal epithelial cells. (A) Phase image of cells isolated from porcine pancreatic duct, showing epithelial-like morphology (bar = 1,000 μm). (B) Immunofluorescent staining for cytokeratin 19 in the cultured primary cells (bar = 200 μm). (C) Immunoblot for the p53 mutant in nine different cells lines transduced with the GKP virus (PGKP cells). Pancreas = wild type pancreatic ductal epithelial cells. Representative blot of three separate experiments. (D-E) qPCR of KRAS and p53 mutants in the nine PKGP cell lines. Each bar represents mean of three separate experiments. *p<0.05, ** p<0.03, ***p<0.01 (unpaired t-test, compared to wild type).

### Generation of tumorigenic cell lines

In order to transform our primary porcine pancreatic ductal epithelial cells, we first generated a lentiviral construct containing *KRAS*^G12D^ and *TP53*^R167H^, genes previously identified [*29*] as the porcine equivalents to the mutant *KRAS* and *TP53* which are present in multiple human cancers [*36–40*]; in the mouse, expression of these mutants was the basis for the KRAS/p53 genetically engineered murine model of pancreatic cancer [*41*]. For our model, we chose to use a lentiviral platform for the vector, because its genome would be large enough to accommodate insertion of both mutant genes; in addition, we believed that a lentivirus would be optimal for transforming primary cells.

Since initial sequencing of the porcine genome has been accomplished [*42*], we were able to utilize the National Library of Medicine’s nucleotide BLAST^®^ database (blast.ncbi.nlm.nih.gov/blast.cgi) to determine the porcine genetic equivalents for human *SMAD4* and *CDKN2A*. We then designed primers (Table S3) to amplify these two genes from genomic DNA isolated from skin of a healthy domestic pig. We sequenced our amplification products, and accessed the BLAST^®^ database to confirm that our products aligned with the porcine *SMAD4* and *CDKN2A* gene sequences. We then proceeded to generate lentiviral constructs to transform primary porcine pancreatic ductal epithelial cells into cell lines expressing various combinations of mutant KRAS and p53, SMAD4 shRNA, and p16^Ink4A^ shRNA (see cell line definitions in Table 1), as described under Materials and Methods.

Primary porcine epithelial cells next were transduced with the GKP lentivirus to generate transformed cell lines. Overexpression of KRAS^G12D^ and p53^R167H^ was confirmed in these cell lines with qPCR and immunoblotting (**Fig. 1C-E**). Of note, we could not obtain a reliable antibody to detect porcine KRAS with immunoblotting, so we had to rely on qPCR results and expression of GFP as markers of KRAS^G12D^ expression. Preliminary *in vitro* analyses to probe the tumorigenic properties of these GKP-transformed cell lines demonstrated modest increases in soft agar colony formation and migration speed over wild type cells (**Fig. 2A & B**). Interestingly, the cell line (2.22) with the best performance in the soft agar and migration assays had only modest overexpression of KRAS^G12D^ and p53^R167H^ (**Fig. 1 & 2**); in contrast, the cell lines with the highest mutant overexpression performed relatively poorly in these *in vitro* assays of “tumorigenesis” (i.e., evidence of transformed behavior in cell culture).

**Fig. 2.**
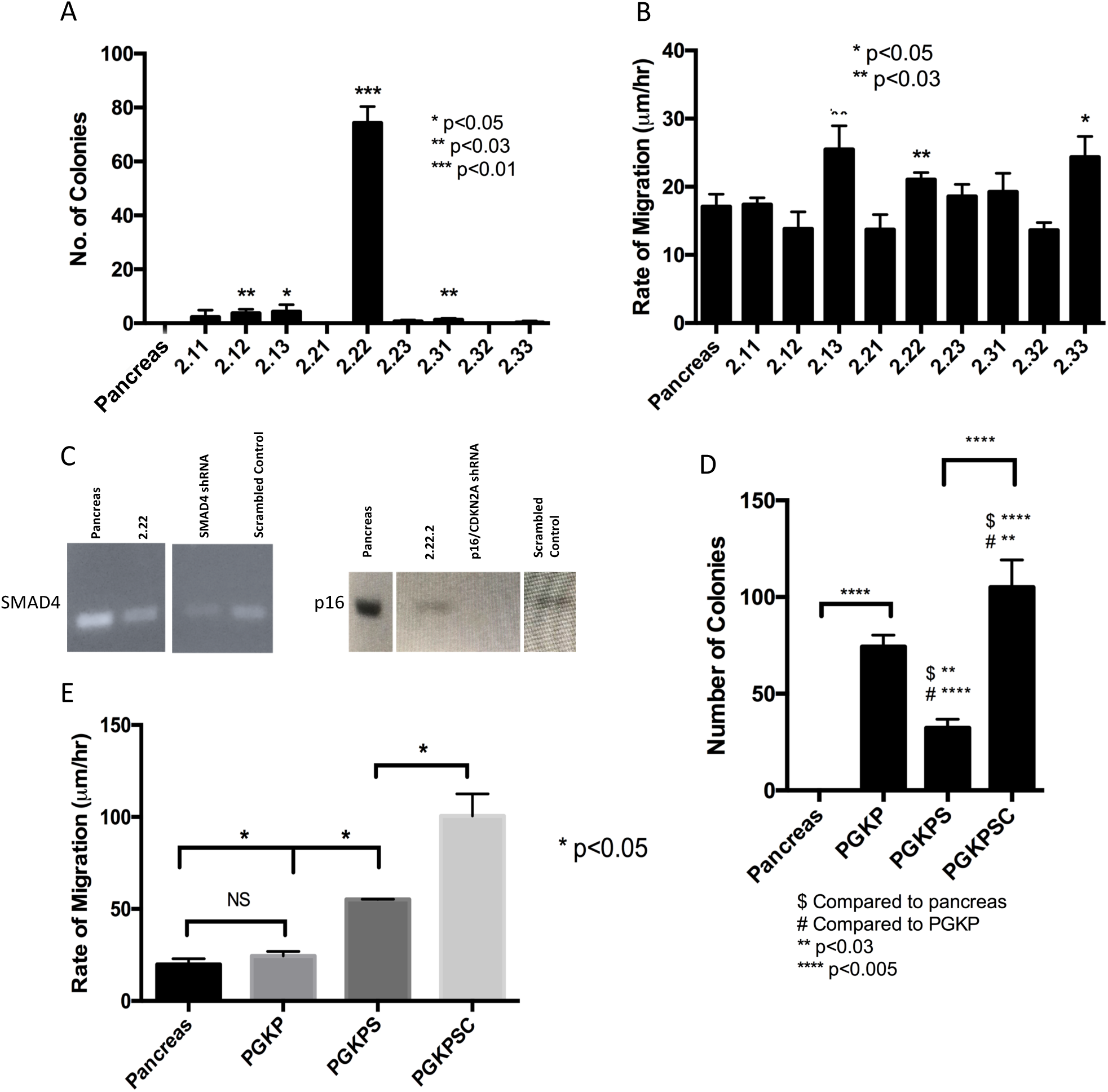
Effect of SMAD4 and p16^Ink4A^ knockdown on transformation in the PGKP cell line. (A) Soft agar assay and (B) migration assay in the nine GKP-transduced cell lines. Pancreas = wild type pancreatic ductal epithelial cells. Each bar represents mean of three separate experiments. (C) RT-PCR of SMAD4 and p16^Ink4A^/CDKN2A mRNA in cell lines expressing targeted or scramble shRNA. Representative blot of three separate experiments. (D) Soft agar assay and (E) migration assay of selected cell lines (PGKP, PGKPS, and PGKPSC), showing the *in vitro* effect of additional knockdown of SMAD4 ± p16^Ink4A^ on the transformation of GKP-transduced pancreatic ductal epithelial cells.

While *in vitro* experiments were being performed, *in vivo* pancreatic tumor induction was attempted utilizing a transgenic mini-pig available from the NSRRC. Known as the “Oncopig,” [*26*], this subject carries an LSL-cassette containing the dominant negative TP53^R167H^ and the activated KRAS^G12D^ sequences [*26,43,44*]; i.e., this subject is the porcine analog of the KRAS/p53 mouse [*41,45*]. As demonstrated previously, site-specific expression of Cre recombinase in the Oncopig resulted in localized p53 inhibition and KRAS activation, while subcutaneous injection of AdCre produced mesenchymal tumors at the injection sites [*26*]. We injected Cre recombinase into the pancreas of five Oncopigs (**Protocol S2; Fig. S2; Tables S5** and **S6**). After four months, we observed no gross tumors. However, there was immunohistochemical evidence of transgene expression at the pancreatic injection sites along with numerous microscopic proliferative lesions with desmoplastic features (**Fig. S2**).

Based on the modest evidence of *in vitro* transformation and the lack of gross *in vivo* tumorigenesis using expression of p53^R167H^ and KRAS^G12D^ only, we decided that additional oncogenic stress might be helpful to increase the tumor-like properties of transformed pancreatic ductal epithelial cells. Utilizing cell line 2.22 (hereafter referred to PGKP; see Table 1), which had relatively good performance in the soft agar and migration assays (Fig. 2A & B), sequential transduction with lentiviral constructs expressing shRNA against SMAD4 and then p16^Ink4A^ was performed to generate cell lines PGKPS and PGKPSC (see Table 1), respectively. RT-PCR then was used to confirm knockdown of the targeted transcripts in these two cell lines (**Fig. 2C**).

### *In vitro* tumorigenic properties of transformed cells

The *in vitro* “tumorigenic” properties of the PGKP, PGKPS, and PGKPSC cell lines first were compared with the soft agar and migration assays (**Fig. 2D & E**). Addition of SMAD4 ± p16^Ink4A^ knockdown enhanced the ability of transformed cells to form colonies in soft agar and increased their migration speed (2D wounding assay), particularly when both transcripts were targeted (i.e., the PGKPSC line). We then compared population doubling time, proliferation (metabolic dye conversion), and Matrigel^®^ invasion ability among the three transformed lines with respect to wild type cells (**Fig. 3A-C**). Both the PGKPS and PGKPSC cell lines had greater proliferation and invasive ability compared to either wild type cells or the PGKP cell line (**Fig. 3B, C**). The doubling time for all three transformed cell lines was approximately the same at ~15 h, compared to the ~4 d doubling time of wild type cells (**Fig. 3A**). Based on the *in vitro* assays of tumorigenesis, we suspected that all three of our transformed cell lines had the potential to form tumors *in vivo*, albeit to varying degrees. We subsequently decided to compare the *in vivo* tumorigenicity among all three cell lines in an immunodeficient mouse model.

**Fig. 3.**
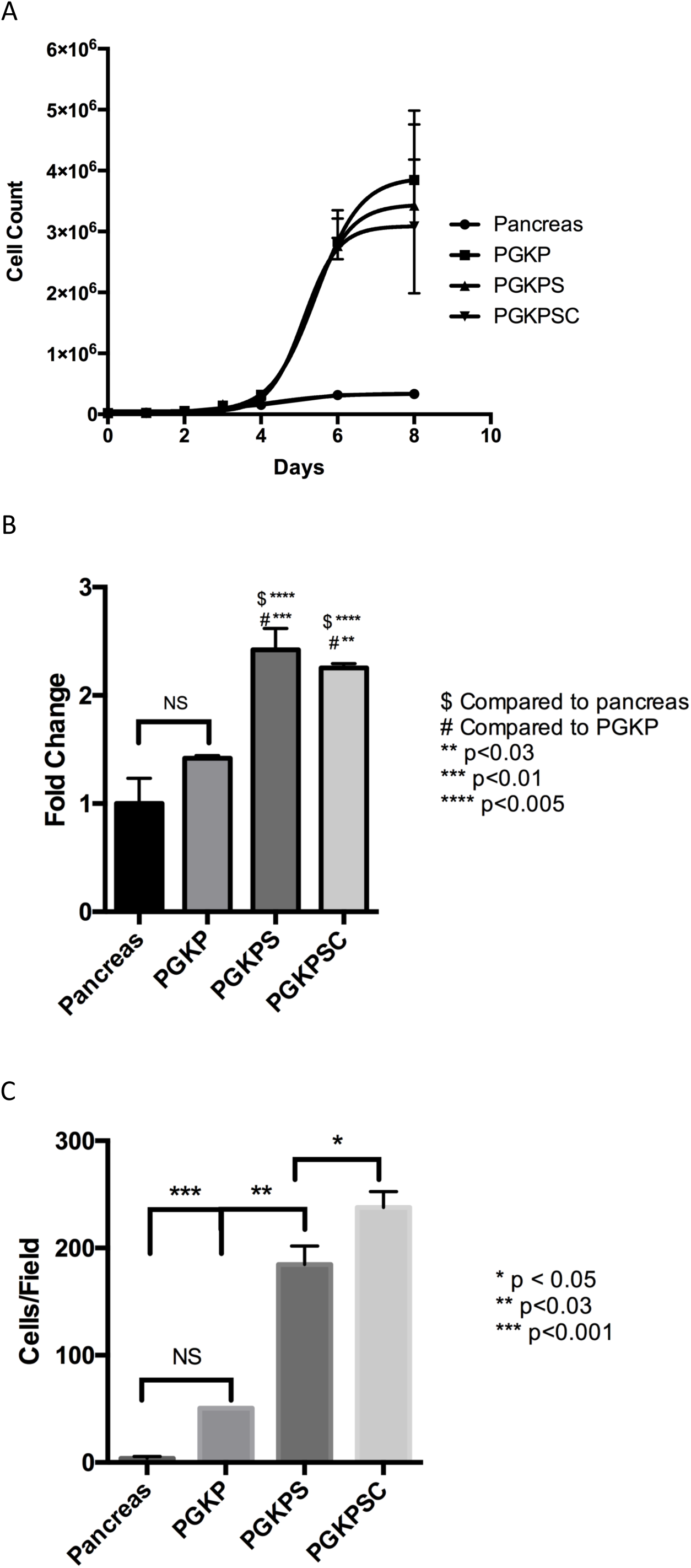
*In vitro* transformation assays comparing the PGKP, PGKPS, and PGKPSC cell lines. (A) Cell culture population doubling time (count-based assay). Pancreas = wild type pancreatic ductal epithelial cells. (B) Proliferation rate (metabolic dye-based assay), represented as fold change, normalized to wild type cells. (C) Invasion (Matrigel^®^-based assay). Each bar or data point represents mean of three separate experiments.

### Subcutaneous and orthotopic tumor transplants in immunodeficient mice

In order to determine if our cell lines retained their tumorigenic properties *in vivo*, we utilized homozygous athymic mice to generate subcutaneous and orthotopic cell implantation models. The subcutaneous model was used to assess *in vivo* tumor growth. All three cell lines grew sizeable tumors (>1 cm diameter) within 6 weeks; growth rates were not statistically different among the three cell lines (**Fig. 4A & B**). The subcutaneous tumors were well-vascularized and mucinous in gross appearance (**Fig. 4A**). Confirmation of GKP lentiviral transduction was demonstrated with immunohistochemistry of the p53^R167H^ mutant in subcutaneous tumors (**Fig. S3**).

**Fig. 4.**
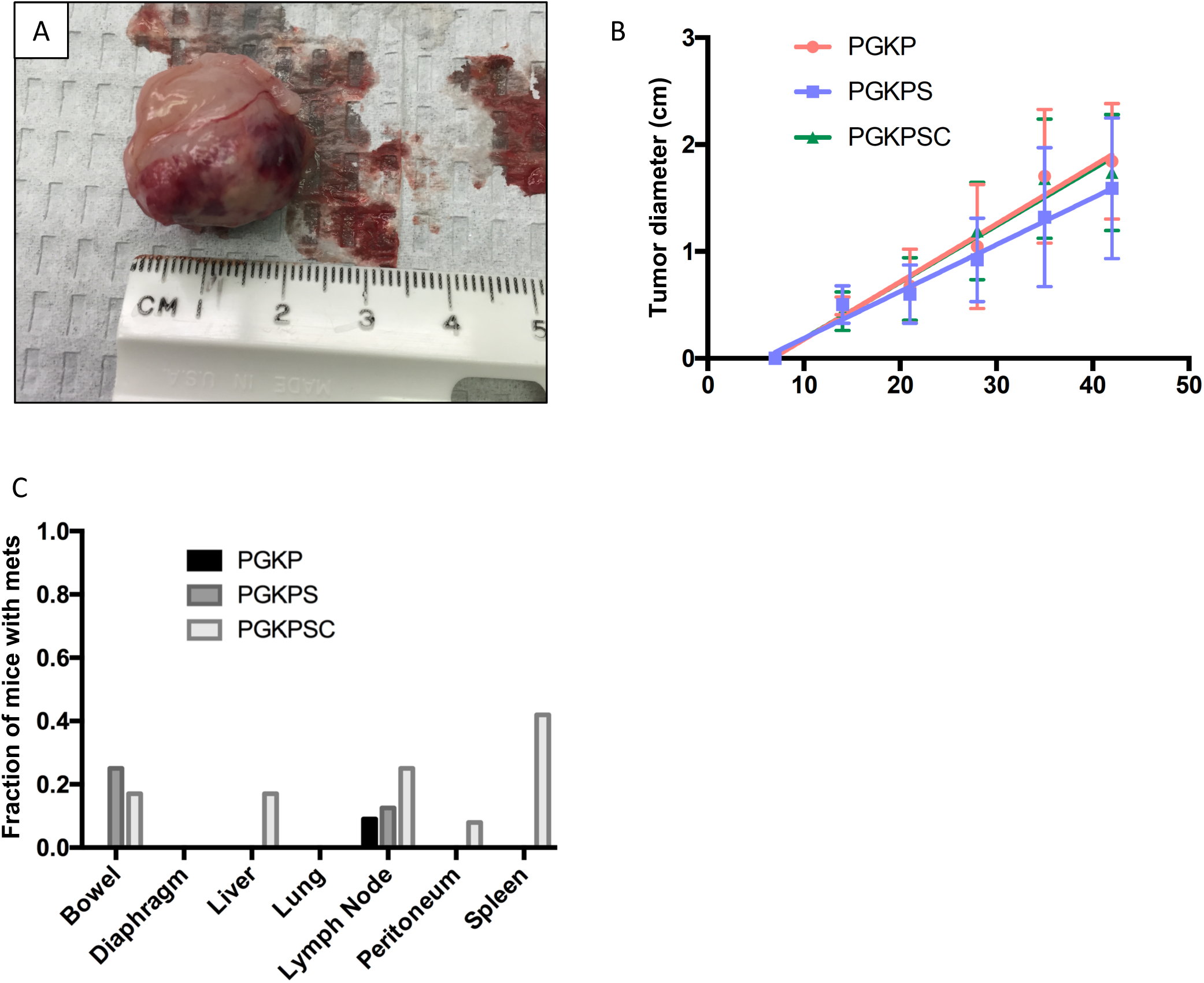
*In vivo* tumorigenesis assays comparing the PGKP, PGKPS, and PGKPSC cell lines. (A) Sample of a resected tumor from subcutaneous injection in nude mice; note size and vascularity. (B) Tumor growth curve from the subcutaneous injection in nude mice. Each data point represents mean of 10 mice. (C) Metastasis after orthotopic implantation in nude mice

Tumors from the subcutaneous implant model then underwent immunohistochemical staining with an array of epithelial and mesenchymal markers (**Fig. 5**). The distribution of staining for the epithelial markers (E-cadherin, epithelial cell adhesion molecule, pan-cytokeratin, cytokeratin-19) generally was more diffuse than the mesenchymal marker staining. In some regions the epithelial marker staining was clustered and intense. The overall abundance of staining for the epithelial markers appeared greater in the PGKPS and PGKPSC lines with respect to the PGKP line. The distribution of staining for the mesenchymal markers (a-smooth muscle actin, vimentin, and type I collagen) was variable, sometimes appearing in cords or strands in some sections, reminiscent of the desmoplastic reaction in human pancreatic cancer [*46,47*]. In other sections, mesenchymal marker staining was minimal. Similar to the epithelial markers, the overall abundance of staining for the mesenchymal markers appeared greater in the PGKPS and PGKPSC lines compared to the PGKP line.

**Fig. 5.**
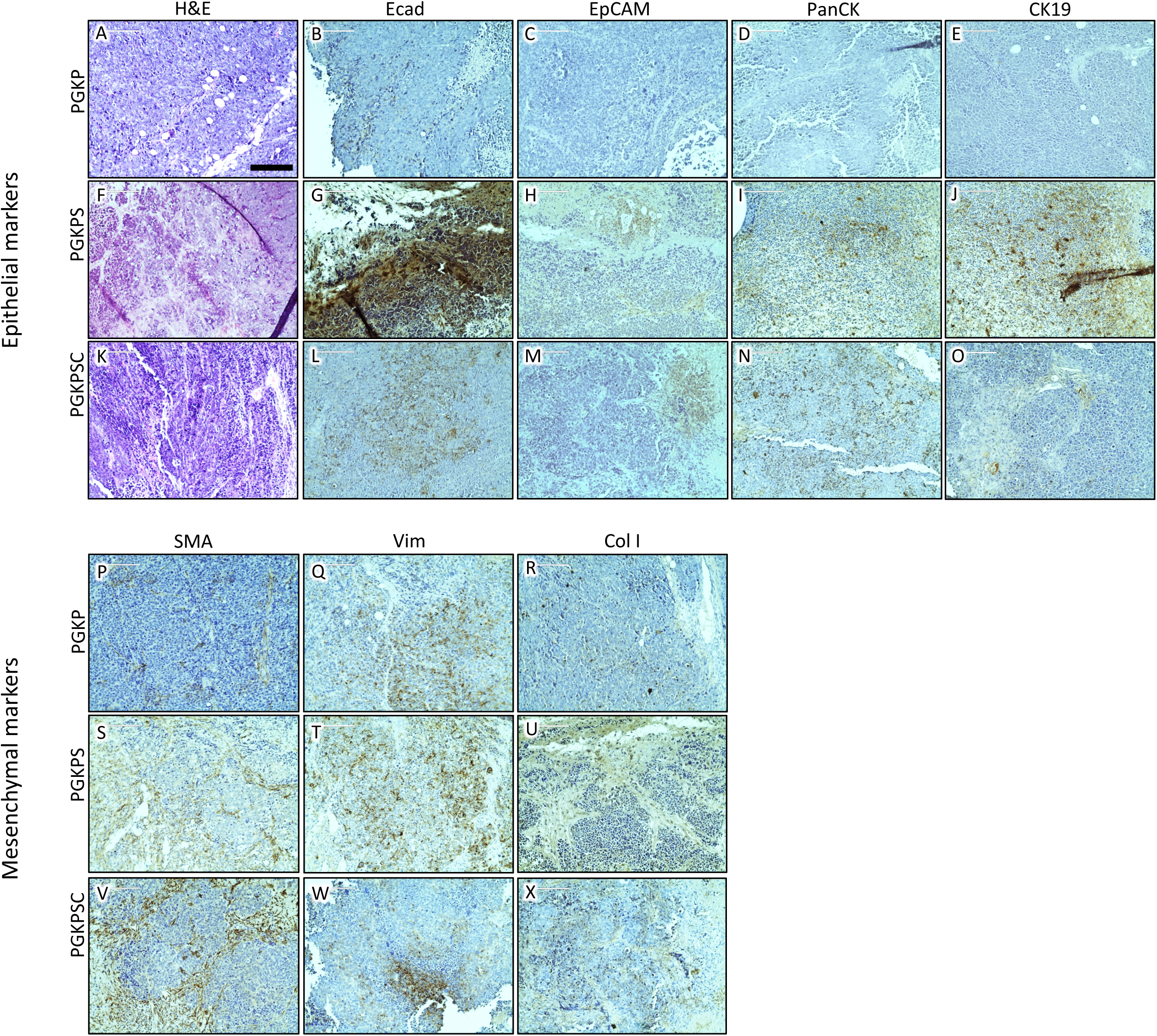
Tumor immunohistochemistry from subcutaneous nude mouse assay. Representative images of primary tumors derived from the PGKP, PGKPS, and PGKPSC cell lines stained with an array of epithelial markers, including E-cadherin (Ecad; B, G, L), Epithelial Cell Adhesion Molecule (EpCAM; C, H, M), Pan-Cytokeratin (PanCK; D, I, N), and Cytokeratin-19 (CK19; E, J, O); and also stained with mesenchymal markers, including a-Smooth Muscle Actin (SMA; P, S, V), Vimentin (Vim; Q, T, W), and Type I Collagen (Col I; R, U, X). Bar = 200 μm.

With the immunohistochemical studies of the subcutaneous tumors demonstrating some epithelial characteristics, the nude mouse orthotopic implantation model was used next to assess the metastatic potential of all three cell lines. The percentage of mice implanted with each cell line that subsequently developed metastasis in the small bowel, diaphragm, liver, lung, lymph node, peritoneum, or spleen is shown in **Fig. 4C**. The degree of metastatic spread was minimal; nodal tissue was the only metastatic site common to all three lines. The PGKPCS cell line exhibited the greatest array of metastatic spread (p < 0.002, Chi-square), with the most common site being the spleen (though it was not clear in two of five mice with splenic disease after PGKPSC implantation whether the spleen was involved with extension from the primary tumor, or from direct seeding, or whether these were true metastases). Interestingly, one of the two subjects with liver metastasis after PGKPSC implantation had primary tumor within the gallbladder, rather than within the pancreas. Similar to the subcutaneous tumor model, most tumors in the orthotopic model were well-vascularized and mucinous in gross appearance.

## Discussion

Our overall goal with this project was to generate tumorigenic pancreatic cell lines that could be used in an immunocompetent porcine model of pancreatic cancer. Current murine models for pancreatic cancer will continue to be helpful, particularly for the study of molecular mechanisms. However, murine models are limited in their ability to replicate human biology and size, so a large animal model of pancreatic cancer likely would enhance our ability to develop and test new diagnostic and treatment modalities for this disease. The data presented herein demonstrated that wild type porcine pancreatic ductal epithelium can be transformed with modulation of common tumor-associated target genes, and that these transformed cells subsequently can grow tumors in immunodeficient mice. These data provide a pathway for the construction of an autochthonous porcine model of pancreatic cancer, namely, orthotopic implantation of tumorigenic pancreatic cells. Proof-of-principle for tumor growth in pigs using subcutaneous implantation of *ex-vivo* transformed autologous fibroblasts was demonstrated in 2007 [*29*].

Porcine biomedical models have been used for decades in the fields of trauma and hemostasis [*48*], xenotransplantation [*49,50*], dermal healing [*51*], toxicology [*52*], atherosclerosis [*53*], and cardiac regeneration [*54*]; the utility of these models is growing. A porcine genome map was generated in 2012 [*42*], and further coverage, annotation, and confirmation is ongoing [*55,56*]. Porcine-centered online tools and databases are now available [*57*]. Genetic manipulation of pigs (including knockouts, tissue-specific transgenics, inducible expression [*29,58–65*]) with similar tools as used in the mouse is becoming more routine, with new gene-edited porcine models emerging in 2015–2017 for diseases such as atherosclerosis, cystic fibrosis, Duchenne muscular dystrophy, and ataxia telangiectasia [*44,66,67*].

The rationale to build a porcine model of pancreatic cancer is (1) to have a platform for diagnostic/therapeutic device development otherwise not achievable in murine models; and (2) to have a highly predictive preclinical model in which anti-cancer therapies (including immunotherapies) could be vetted/optimized prior to a clinical trial [*68*]. The rationale to use the pig in this modeling effort is that this species mimics human genomics [*55,69–72*], epigenetics [*73*], physiology [*52,69,74*,75], metabolism [*69,75,76*], inflammation and immune response [*72,77–81*], and size [*75,82*] remarkably well (in particular, better than mice), with reasonable compromises towards cost and husbandry [*75*]. So based on the pig’s relatively large size and its proven track record in replicating human biology which, incidentally, is a demonstrably better replication than can be obtained with rodents, we selected swine as the model organism for this pancreatic cancer project.

Research on immunocompetent large animal cancer models [*83–85*] includes prostate cancer, for which there is a canine model [*86*]. In addition, in 2012 a group in Munich reported the engineering of (i) an *APC* mutant pig that developed rectal polyposis [*16,57*] and (ii) a pig with Cre-inducible p53 deficiency [*63*]. This group subsequently determined that their p53-null subjects (*TP53*^R167H/R167H^) developed osteosarcoma by age 7–8 months [*88*]. Other p53-deficient pigs have been engineered since this initial report [*64,89*]; in the report from Iowa, half (5 out of 10) of p53-deficient (*TP53*^R167H/R167H^) pigs developed lymphoma or osteogenic tumor at age 6–18 months [*64*]. A group in Denmark reported the creation of a *BRCA* mutant pig in 2012. [*90*].

A KRAS/p53 “Oncopig” was reported in 2015 [*26,43,44,84*]. This subject has a somatic LSL-cassette that can express dominant negative p53 (R167H mutation) and activated KRAS (G12D mutation); i.e., the porcine analog of the KRAS/p53 mouse [*41,45*]. Site-specific expression of Cre recombinase in the Oncopig resulted in localized p53 inhibition and KRAS activation; subcutaneous injection of AdCre produced mesenchymal tumors at the injection sites [*26*]. In 2017, initial work was published on a Oncopig-based model of hepatocellular carcinoma [*91*]. Also in 2017, another genetic porcine model of intestinal neoplasia was reported [*92*], utilizing inducible expression of KRAS^G12D^, c-Myc, SV40 large T antigen, and retinoblastoma protein (pRb). One out of three pigs total in this model developed duodenal neuroendocrine carcinoma with lymph node metastasis at two months after induction. A porcine model of pancreatic cancer has not yet been reported, other than some preliminary data presented by us in 2017 [*93*].

The process we used for developing transformed porcine pancreatic ductal epithelial cell lines for future orthotopic implantation was somewhat iterative, in that we modified our strategy along the way based on our early results. Commonly mutated genes in pancreatic cancer include *KRAS* [*94,95*] and *TP53* [*95–97*]. In mice, somatic activation of KRAS via the G12D mutation (KRAS^G12D^) produced widely metastatic pancreatic tumors; survival duration in these subjects decreased further with p53 inactivation [*41*]. Based on this murine model, and the published success with subcutaneous tumor induction in the KRAS/p53 Oncopig [*26*], we elected to transform pancreatic ductal epithelial cells with expression of activated KRAS^G12D^ and p53^R167H^ only. However, the initial results from our *in vitro* transformation assays with these two gene edits (i.e., the PGKP cell line of Table 1) were somewhat underwhelming. Combined with the finding of no gross tumor four months after pancreatic AdCre injection in five Oncopig subjects, we decided that additional genetic “hits” might be necessary for transformation of porcine pancreatic ductal epithelial cells. Of note, Schook et al. [*29*] found that porcine dermal fibroblasts required six genetic edits (human telomerase reverse transcriptase, dominant negative p53, cyclin D1, activated cyclin dependent kinase, oncogenic c-Myc, and oncogenic H-Ras) to optimize the tumorigenic phenotype of this particular cell.

Other commonly mutated genes in pancreatic cancer include *SMAD4* [*95,98*] and *CDKN2A* [*95,99*]. Deletion of *SMAD4* or *CDKN2A* in a *KRAS*^G12D^ murine pancreatic cancer model enhanced tumor growth [*100,101*]. Based on these published data and our above transformation results with just the *KRAS* and *TP53* edits, we elected to add knockdown of SMAD4 and p16 to our list of hits for transformation of porcine pancreatic ductal epithelial cells. Ultimately, all three of our cell lines (Table 1) demonstrated transformed behavior *in vitro* and the ability to form tumors *in vivo* (nude mice), with perhaps some enhancement by the addition of SMAD4 and p16 knockdown. In the future, we intend to utilize CRISPR/Cas9 editing to disrupt the genes of these and/or other targets.

Although we utilized a relatively high number of cells to obtain tumor formation in our nude mice experiments, we feel that this is due to the relative young age of our cell lines, which did not undergo as many passages as other cell lines which have been used to study pancreatic cancer. Thus, the transformed cell lines in this report likely did not undergo an inadvertent selection for the fastest growing cells, as may occur in older, extensively-passaged lines. Regarding the lack of desmoplasia in our murine xenograft model, we did not consider this surprising, in that most immunodeficient murine models of pancreatic cancer do not recapitulate the extent of desmoplasia or metastasis seen with human disease. We anticipate that future experiments involving the implantation of transformed pancreatic ductal epithelial cells into wild type swine (and thus into a “normal” microenvironment with “normal” immunoediting) will produce tumors with desmoplasia and metastasis.

Of note, the pathologic findings in our five Oncopig subjects, while not macroscopically tumorous, displayed tissue architecture reminiscent of desmoplasia. Regarding the lack of macroscopic tumor in these transgenic pigs, it is conceivable that the induction process was not optimal secondary to an inadequate dose of AdCre, inadequate recombinase activity, inadequate tissue delivery of the enzyme, or some other technical issue. Our future plans in this respect will involve additional attempts at induction of pancreatic tumor in the Oncopig with and without introduction of some additional gene edits, such as local disruption of *SMAD4* and *CDKN2A* (e.g., with *in vivo* CRISPR/Cas9 gene editing [*102,103*]).

## Acknowledgements

This study is the result of work supported in part with resources and the use of facilities at the Omaha VA Medical Center (Nebraska-Western Iowa Health Care System). Portions of this study were presented at the 108th Annual Meeting of the American Association for Cancer Research (AACR) in Washington, D.C., April 1–5, 2017. The authors would like to acknowledge the technical assistance of Gerri Siford, Chris Hansen, Kelly O’Connell, and Tom Caffrey. The authors also would like to thank Dr. Lawrence B. Schook and Dr. Laurie A. Rund at the University of Illinois for the gift of their plasmid that contained the *KRAS*^G12D^ mutation, along with comments, insights and suggestions for its use in this project. Datasets from this manuscript will be freely shared upon request to the senior author (MAC).

## Author Contributions

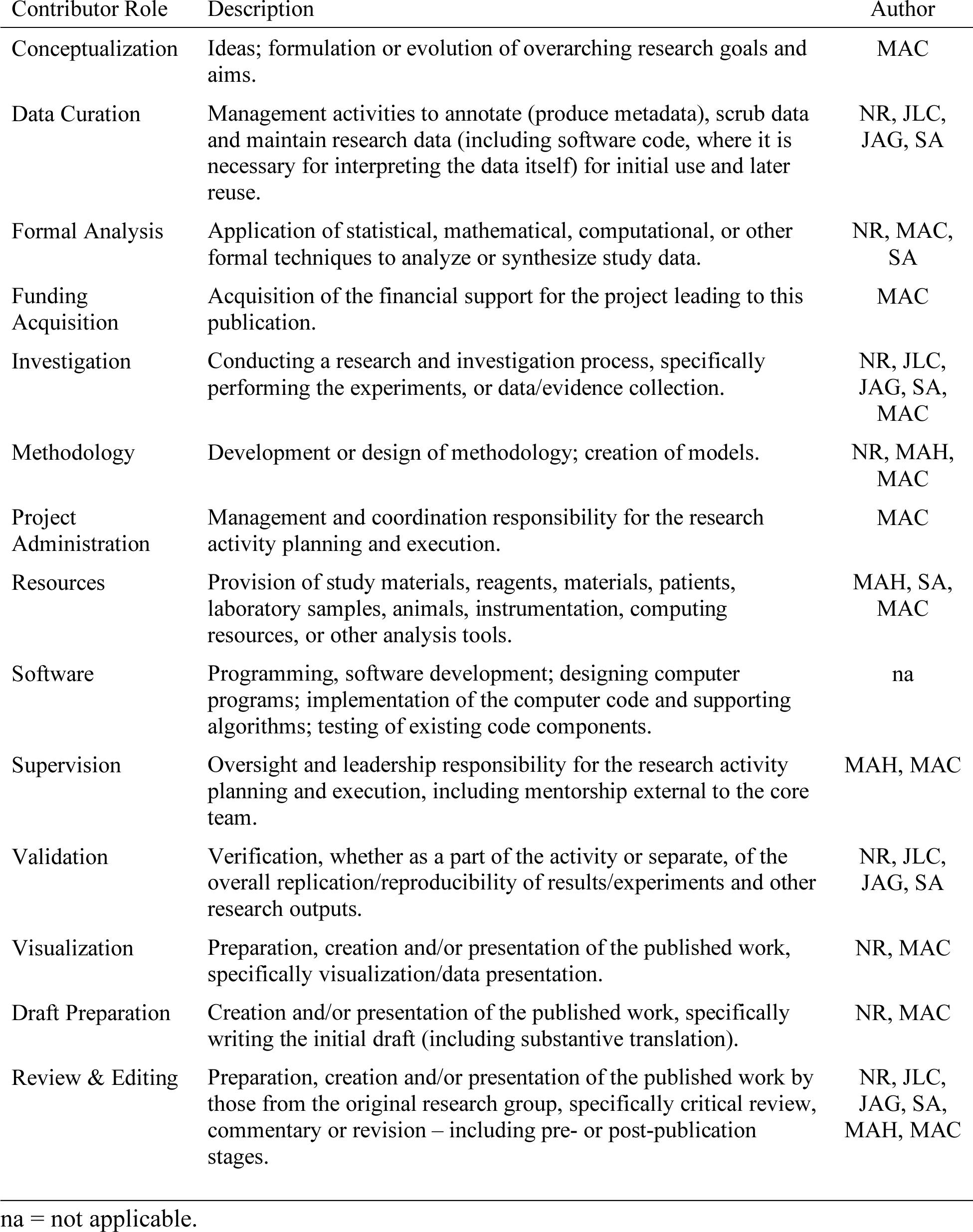

## Supporting Information

**Fig. S1.** pIRES2-AcGFP1 Vector Information

**Fig. S2.** Studies with the NSRRC KRAS/p53 Oncopig

**Fig. S3.** KRAS/p53 immunohistochemistry of subcutaneous murine tumors

**Protocol S1.** Isolation of Epithelial Cells from Porcine Pancreas

**Protocol S2.** Porcine methodology

**Table S1.** Responses to the ARRIVE recommendations

**Table S2.** Responses to the NIH Preclinical Research Guidelines

**Table S3.** Primers and other short sequences

**Table S4.** Antibody information

**Table S5.** Oncopig descriptive data

**Table S6.** Oncopig serum laboratory testing

